# Retroviral infection and commensal bacteria dependently alter the metabolomic profile in a sterile organ

**DOI:** 10.1101/2023.01.10.523497

**Authors:** Jessica Spring, Vera Beilinson, Brian C. DeFelice, Juan M. Sanchez, Michael Fischbach, Alexander Chervonsky, Tatyana Golovkina

## Abstract

Both viruses and bacteria produce ‘pathogen associated molecular patterns’ that may affect microbial pathogenesis and anti-microbial responses. Additionally, bacteria produce metabolites while viruses could change metabolic profiles of the infected cells. Here, we used an unbiased metabolomics approach to profile metabolites in spleens and blood of Murine Leukemia Virus-infected mice monocolonized with *Lactobacillus murinus* to show that viral infection significantly changes the metabolite profile of monocolonized mice. We hypothesize that these changes could contribute to viral pathogenesis or to the host response against the virus and thus, open a new avenue for future investigations.

## Introduction

The intestinal commensal microbiota is a key factor that mediates host health by providing nutrients and vitamins, supporting the development and shaping of the secondary lymphoid organs in the intestine, and conferring resistance of colonization by pathogenic microorganisms (1). Many of these, and other microbiota dependent processes, are mediated by a number of commensal bacteria derived high- and low-abundance molecules. Various commensal small molecules exhibit potent biological activities that can influence the host’s cellular activities (2-4). Of these small molecules, metabolites have been shown to serve as an energy source, modulate diseases and psychiatric disorders, promote intestinal barrier function, and modulate the immune system (5-7)

Metabolites have both positive and negative effect on pathogenic bacteria. These effects include enhanced biofilm formation, increased expression of virulence factors (8-10), disruption of bacterial cell structures, suppression of bacterial growth, and stimulation of innate immune cell proliferation (8, 11-14). Viral infections are also impacted by metabolites. Metabolites can be used as energy sources to promote viral replication (15, 16). Whereas other metabolites promote the immune response and augment interferon (IFN) expression in a variety of viral infection models (17, 18).

While the effect of commensal microbe-derived metabolites on cellular activities and pathogen infections is clear, a reciprocal impact of the commensal microbiota and viral pathogens on metabolites has yet to be investigated. We previously showed commensal bacteria *Lactobacillus murinus* promotes the pathogenesis of murine retrovirus Murine Leukemia Virus (MuLV) (19). We used a similar model coupled with an unbiased metabolomics approach to investigate the influence of cross-talk between commensal bacteria and viral infection on metabolites. Here, we show viral infection and commensal bacterial colonization dependently alter the metabolomic profile in a sterile organ.

## Methods

### Mice

Breeding and maintenance of mice used in this study was conducted at the animal facility of The University of Chicago. BALB/cJ mice were bought from The Jackson Laboratory (TJL). Females and males were used at a ratio of ∼50:50. The Animal Care and Use Committee at The University of Chicago reviewed and approved the studies described here.

### Monitoring sterility in GF isolators

BALB/cJ mice were re-derived as germ-free (GF) at Taconic farms and subsequently housed in sterile isolators at the gnotobiotic facility at the University of Chicago. GF isolator sterility was determined as previously described (20). Fecal pellets from isolators were collected weekly and quickly frozen. A bead-beating/phenol-chloroform protocol was used to extract DNA. DNA was amplified using primers that widely hybridize to bacterial 16S rRNA gene sequences. In addition, microbiological cultures were established with GF fecal pellets, specific pathogen free (SPF) fecal pellets (positive control), sterile saline (sham), and sterile culture medium (negative control). Samples were inoculated into BHI, Nutrient, and Sabbaroud Broth media and incubated aerobically and anaerobically at 37°C and 42°C. Cultures were maintained for five days until deemed negative.

### Colonization of GF mice

*Lactobacillus murinus* (*L. murinus* ASF361) was isolated from the Altered Schaedler Flora (ASF) consortium (21). Accordingly, fecal matter from ASF colonized gnotobiotic mice was suspended in sterile PBS and plated onto a selective media for *Lactobacilli*, de Man, Rogosa and Sharpe agar (MRS) (ThermoFisher Scientific). The identity of *L. murinus* was confirmed by sequencing of the 16S RNA ribosomal amplicon generated via PCR from a bacteria colony. GF BALB/cJ mice were monocolonized with *L. murinus* by oral gavage of 200µl of overnight liquid culture grown from a single colony. Monocolonization was validated by sequencing PCR products generated from fecal DNA and 16S rRNA primers and was verified at closing of the experiment.

### Virus isolation and infection

Rauscher-like Murine Leukemia Virus (RL-MuLV) is composed of N, B tropic ecotropic and mink cell focus forming (MCF) virus (22). RL-MuLV was isolated from tissue culture supernatant of chronically infected SC-1 cells (ATCC CRL-1404). Titers of ecotropic virus within RL-MuLV mixture were determined via an XC plaque assay (23). 1 × 10^3^ pfus were intraperitoneally (i.p.) injected into 6-8 week-old BALB/cJ female mice (G0 mice). G0 mice were bred to produce the progeny G1 mice. Spleens from G1 mice were used as a source of virus. Homogenized spleens from 2-3-month-old G1 mice were centrifuged at 4°C for 15 minutes at 2,000 rpm. Supernatant was collected and layered onto a 30% PBS/sucrose cushion and spun at 31,000 rpm for 1 hour at 4°C in a TW55.1 bucket rotor. The pellet fraction was re-suspended in PBS. Insoluble material was removed by spinning the resuspended fraction at 4°C at 10,000 rpm. Supernatant was collected and aliquoted, titered via plaque assay, and stored at -80°C.

RL-MuLV isolated from spleens was diluted in sterile PBS followed by filtering through a sterile 0.22μm membrane in a laminar flow hood. GF and *L. murinus-*monocolonized BALB/cJ females were injected i.p. with 1 × 10^3^ pfus (G0 mice). G0 females were bred to produce G1 mice. Uninfected and infected GF and *L. murinus*-monocolonized were sacrificed at 2 months. Plasma and spleens were isolated and stored at -80°C. Mice were confirmed to be infected via PCR using primers specific for the LTR of ecotropic virus. Forward primer - 5’ATGAACGACCCCACCAAGT3’ and reverse primer – 5’GAGACCCTCCCAAGGAACAG3’. Spleen weights ranged from: 0.07 to 0.09g among uninfected GF mice; 0.07-0.09g among uninfected *L. murinus-*monocolonized and 0.15 – 0.19g among infected *L. murinus-*monocolonized. Infected mice were non-leukemic as evaluated by histology of H&E stained spleen sections.

### FITC permeability assay

Permeability of the mouse gut was assessed using a FITC permeability assay. Food and water were removed from mouse cages for four hours. After four hours, mice were gavaged with 60mg of FITC-dextran (MW 4,000, Sigma-Aldrich) per 100g of mouse (24). Three hours post gavage, mice were bled into 100µl heparin, and plasma was separated by spinning the blood at 2,000rpm for ten minutes at 4°C. 50µl of each sample was loaded into a 96-well plate in duplicate to measure FITC concentration at emission and excitation wavelengths of 520nm and 490nm, respectively using a TECAN fluorescence spectrophotometer and Magellan software. FITC-dextran standards were diluted in plasma from unmanipulated mice. Fluorescence from the negative control samples (plasma from unmanipulated mice) was subtracted from fluorescence of the standards and experimental samples.

### Metabolomics

Spleens and sera were harvested and maintained at -80°C prior to metabolite extraction. About 10mg per sample was used for the following analysis. Extraction solvent (1:2:2 water:acetonitrile:methanol containing stable isotope labeled internal standards) was added to each spleen sample at a ratio of 400µL per 10mg of tissue. Four to six 2.3mm stainless steel beads were added to each sample tube and method blanks. Samples were homogenized by bead beating at 20 Hz for 15 minutes. Homogenized samples were placed at -20°C for 1 hour to maximize protein precipitation followed by brief vortexing and centrifugation for 5 minutes at 4°C and 14,000 X g. 120 µL of supernatant was taken and passed through a 0.2 µm polyvinylidene fluoride centrifugal filter at 4°C for 3 min at 6,000 X g. Collected extract was stored at 4°C during analysis.

Metabolites were extracted from sera samples as follows. Frozen specimens were thawed on ice and inverted 5 times to mix. From each specimen, 50 µL of serum was aliquoted into new 1.5 mL Eppendorf tubes. 150 µL of 1:1 acetonitrile:methanol, containing stable isotope labeled internal standards, was added to each 50 µL aliquot followed by vortexing for 20 seconds. All tubes were stored at -20°C for 1 hour then promptly vortexed for 20 seconds and centrifuged at -10°C for 10 minutes at 21,130 X g. Supernatant was transferred to a new 1.5 mL Eppendorf tube and dried at room temperature in a LabConco Speedvac. Dried extracts were resuspended in 50 µL 4:1 acetonitrile:water and stored at 4°C during analysis.

Extracted metabolites from both sera and spleen were subjected to hydrophilic interaction liquid chromatography (HILIC) untargeted metabolomics at the Chan Zuckerburg Biohub. Data processing, chromatography, and tandem mass spectral data collection methods have been previously described elsewhere (25).

Metabolites which frequencies had a relative standard deviation (RSD) greater than 30% between samples in the same group were discarded. Metabolites with RSDs less than 30% were normalized with Log_10_, quantile normalization, followed by determining the z-score.

## Results

### Bacterial colonization and viral infection drastically alter the peripheral metabolite landscape

To identify metabolites whose presence and abundance were dictated by both colonization and viral infection, an unbiased metabolomics approach was undertaken. Accordingly, we compared metabolites within the sera and spleens of Murine Leukemia Virus (MuLV) infected and uninfected BALB/cJ mice that were either germ-free (GF) or monocolonized with mouse commensal *Lactobacillus murinus (L. murinus)*. We used Rauscher-like MuLV variant which is spread via blood and replicates within the proliferating erythroid progenitor cells subsequently causing erythroid leukemia (22). GF and *L. Murinus*-colonized ex-GF mice were injected with 0.22µM sterilized virus and bred to produce offspring (G1 mice) infected via a natural route. Sera and spleens from four groups of 2-month-old G1 mice: uninfected GF, ex-GF *L. murinus* colonized, MuLV-infected GF, and MuLV-infected ex-GF *L. murinus* were subjected to coupled liquid-chromatography/tandem-mass spectrometry (LC-MS/MS). Mass spectrometry spectra were collected under positive and negative ionization modes. Metabolites which frequencies had a relative standard deviation (RSD) greater than 30% between samples within the same group were discarded, and the remaining metabolites were subjected to further analyses (Figures 1A-C).

**Figure 1:**
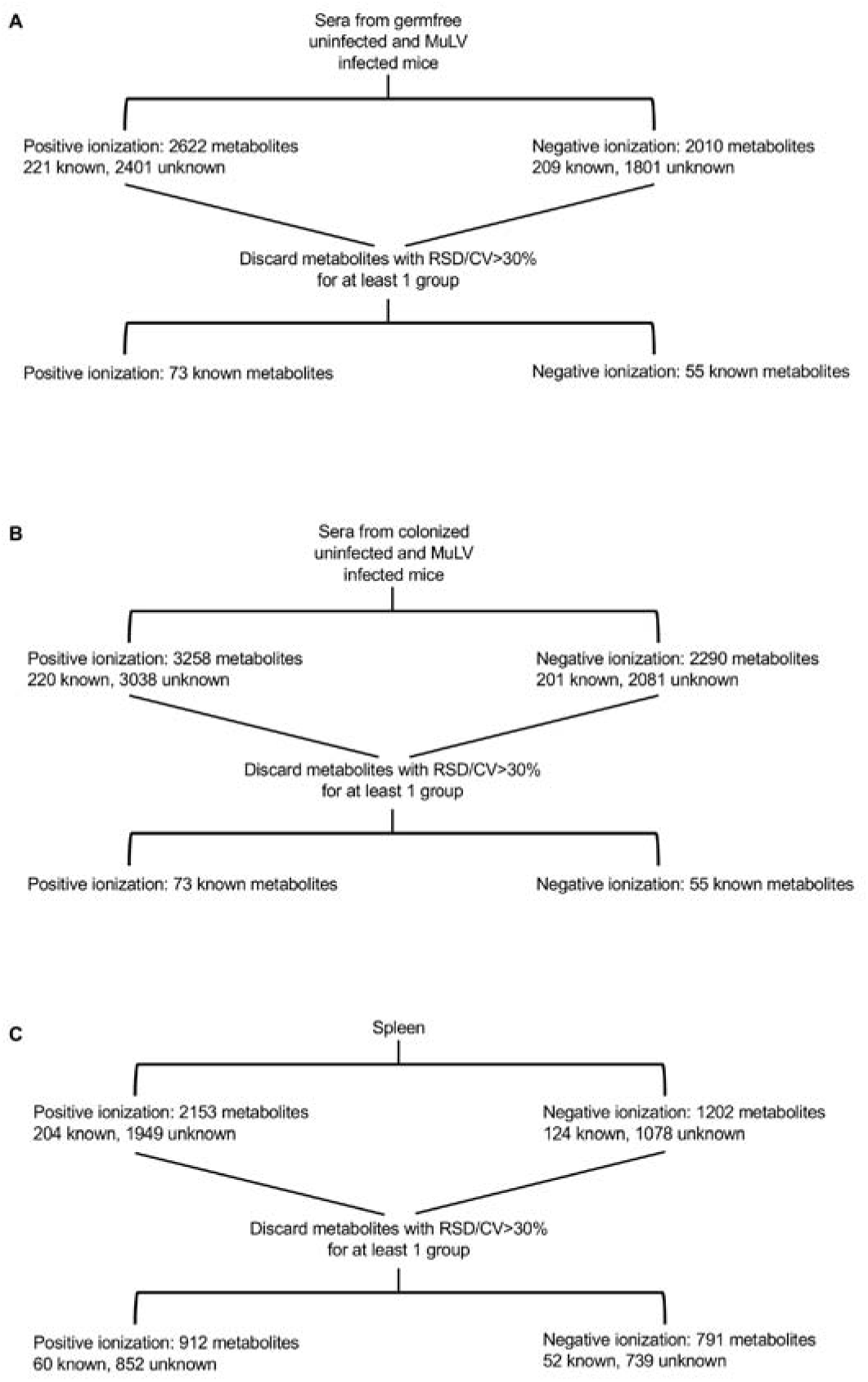
Flow chart illustrating the processing of sera and spleen samples for metabolomics. Mass spectrometry for sera samples was conducted in two separate procedures **(A, B)**. Therefore, unknown metabolites could not be matched between the experiments and were removed. Spleen **(C)** samples were subjected to mass spectrometry under positive and negative ionization modes. Metabolites within a cohort of mice with a relative standard deviation (RSD) > 30% were discarded leaving the indicated number of known and unknown metabolites.

Sera samples collected from uninfected and infected germfree mice (Figure 1A) were subjected to LC-MS/MS separately from sera samples of uninfected and infected colonized mice (Figure 1B), therefore it was impossible to match unknown metabolites between these two groups. Consequently, unknown metabolites from the sera were discarded in later comparative analyses. Only identifiable metabolites found between all four groups of mice were kept for subsequent analyses. This includes 73 and 55 known metabolites identified under positive and negative ionization, respectively (Figures 1A, 1B and Tables S1 and S2). MuLV infection of GF mice marginally altered the overall metabolomic profile (Figure 2A) and the level of metabolites within the sera (Figure 2B). In contrast, *L. murinus* colonization led to the greatest shift in metabolites (Figures 2A and 2B). MuLV infection of *L. murinus* colonized mice induced further minimal changes in the metabolomic landscape and quantity of metabolites within the sera (Figures 2A and 2B).

**Figure 2:**
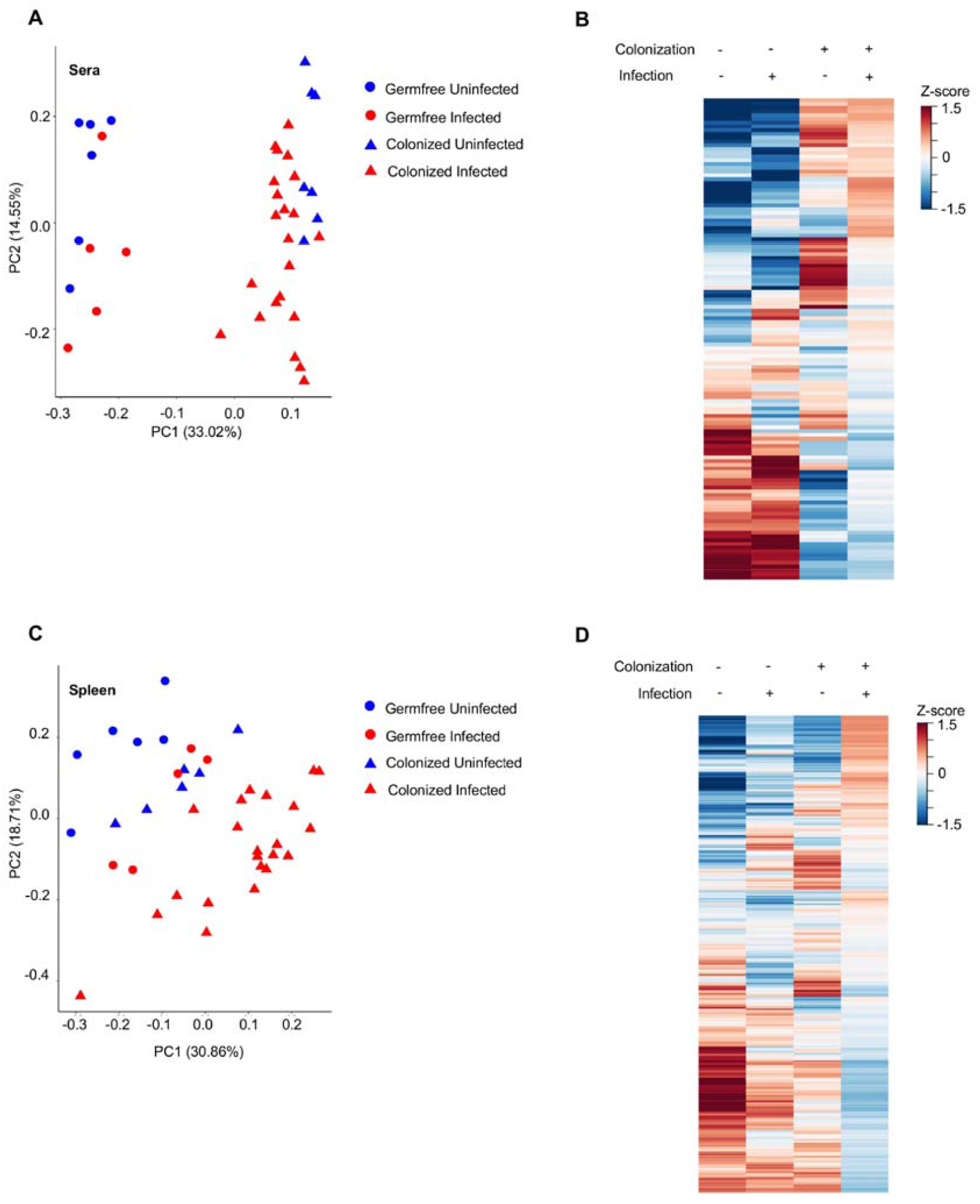
MuLV infection and bacteria colonization alter the metabolomic profile in the sera and spleen. (A, C) PCoA plot of all metabolites within the sera (A) or spleen (C) of the indicated mice. Each point is an individual mouse. (B, D) Heatmaps of metabolite abundances in the sera (B) or spleen (D) of the four groups of mice. Each row is a metabolite.

From the spleens of all mouse groups, a total of 60 known metabolites were identified under positive ionization and 52 under negative ionization (Figure 1C and Tables S2 and S3). Similar to sera, the vast majority of changes were found among over 1,500 unknown metabolites (Figure 1C). Contrary to the sera, splenic metabolites were altered most drastically in the presence of both *L. murinus* colonization and viral infection compared to either condition alone (Figures 2C and 2D). Therefore, viral infection in monocolonized mice led to negligible shifts in the metabolomic profile within the sera, but greatly influenced metabolites within the spleen. These data indicate that bacterial colonization of the gut together with MuLV infection promotes metabolite accumulation or production in the spleen.

Through our analysis, we identified three groups of metabolites which, compared to GF mice, were altered upon MuLV infection or *L. murinus* colonization or both infection and colonization. Group 1 includes metabolites which were significantly altered in *L. murinus-*colonized mice; group 2 includes metabolites which were significantly altered in virally-infected mice; and group 3 consists of metabolites which were significantly altered only in colonized and infected mice. As we were interested in metabolites whose presence and quantity were mediated by colonization and viral infection, we further investigated metabolites identified in group 3. Considering that bacterial colonization alone conferred the largest shift in the metabolomic landscape within the sera (Figure 3A), it was not surprising that only 29 identifiable metabolites were found to be significantly altered in the sera of colonized and infected mice compared to the sera from the other three groups of mice (Figure 3A). In contrast, 271 known and unknown metabolites were found to be impacted by both bacterial colonization and viral infection in the spleen (Figures 3C and 3D). However, the vast majority of them were unknown and only 21 metabolites were identifiable (Figure 3D).

**Figure 3:**
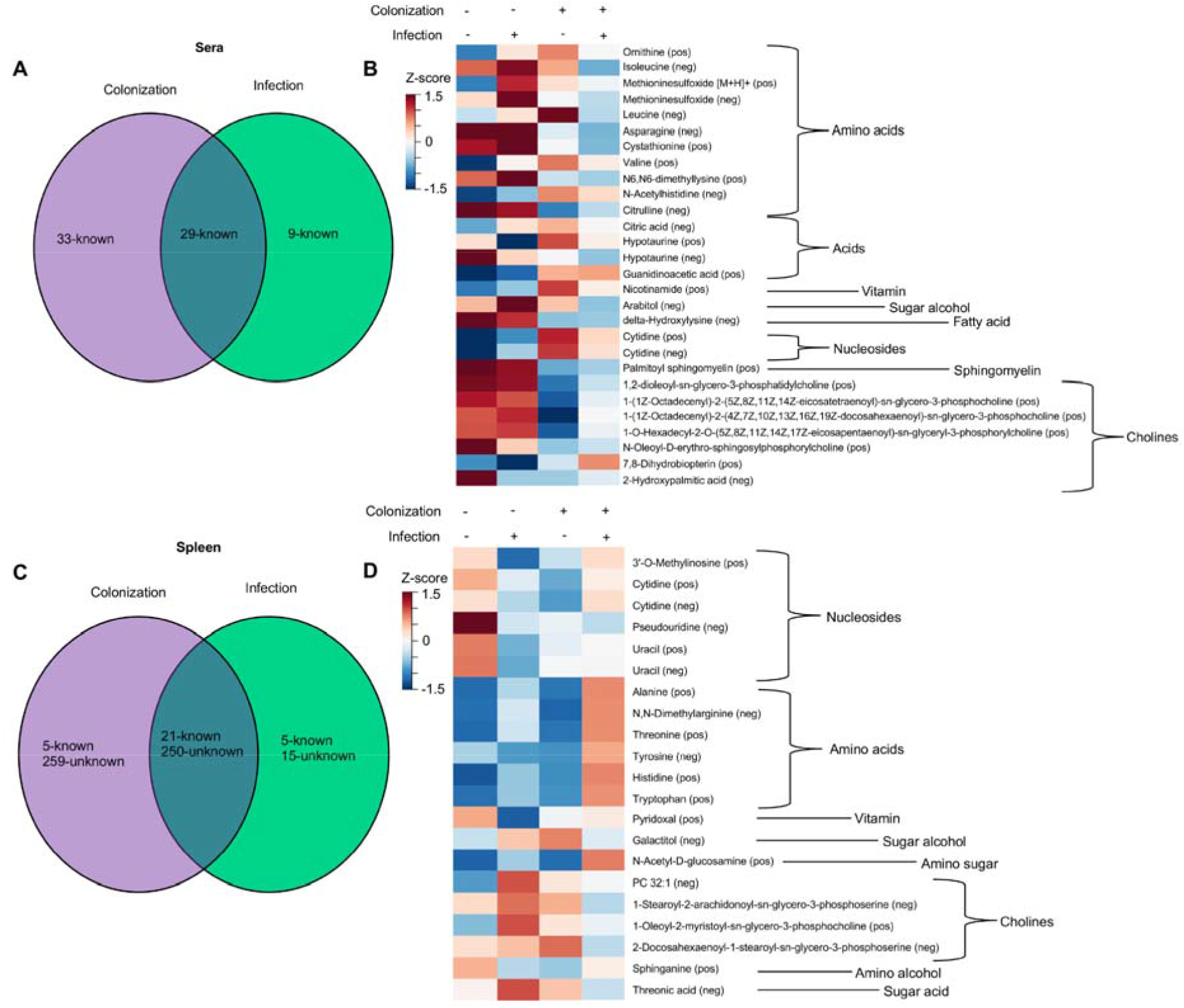
Abundances of certain metabolites is altered by colonization and MuLV infection. (A, C) Venn-diagrams illustrating the numbers of metabolites significantly altered by colonization, infection, or both in the sera (A) or spleen (C). (B, D) Heatmaps of identified metabolites, clustered by chemical type, whose abundances are influence by colonization, MuLV infection, or both in the sera (B) or spleen (D). Normalized values for each group were averaged and constructed into a heat map where red color indicates high metabolite levels and blue color indicates lower metabolite levels. P-values were adjusted using Bonferroni correction (35) followed by two-way ANOVA to identify metabolites influenced by two independent variables: the presence of *L. murinus* and the virus. Each row is a metabolite.

Viral infection has been shown to alter the intestinal barrier, enabling microbial translocation (26, 27). Based on the observation that MuLV infection of colonized mice greatly enhanced the abundance of metabolites within the spleen, we sought to determine whether MuLV increased the permeability of the intestines. To address this possibility, intestinal permeability was compared between uninfected and infected conventionally housed specific pathogen free (SPF) mice. Mice were orally gavaged with MW 4,000 FITC-dextran, a large molecule that will only cross from the intestine into the periphery if the intestinal barrier has been compromised, and fluorescence within the plasma was assayed four hours later. Concentration of FITC detected within the plasma was similar between uninfected and infected SPF mice, suggesting MuLV does not alter intestinal permeability (Figure 4). Thus, enhanced permeability of the intestinal barrier is not the cause of increased metabolites within the sera and spleen in infected mice.

**Figure 4:**
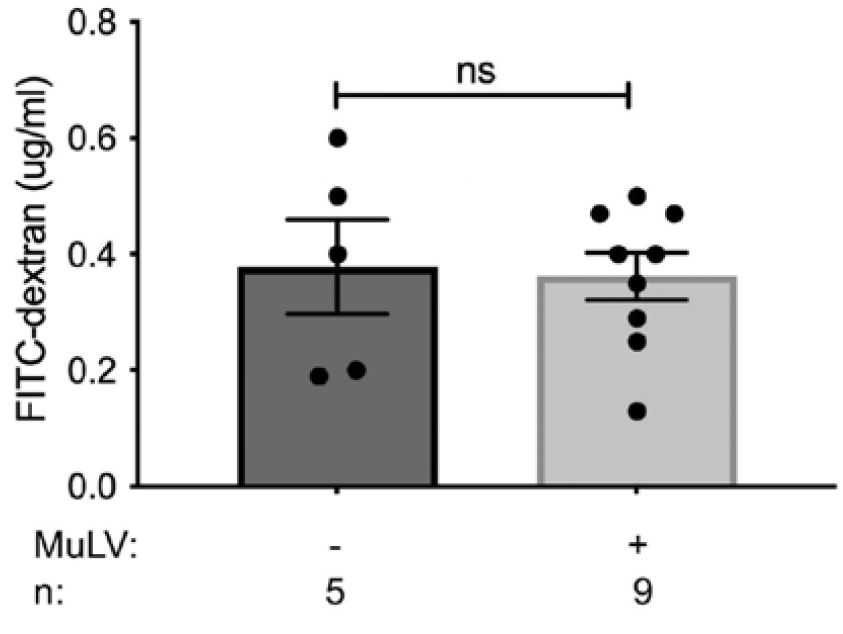
Intestinal permeability is unaffected by MuLV infection. FITC permeability assay was used to determine the permeability of the intestines in SPF BALB/cJ infected and uninfected mice. n, number of mice used. *p* value calculated using unpaired *t* test.

## Discussion

The commensal microbiota has been demonstrated to modulate replication and pathogenesis of viruses from various families (28-31). Furthermore, commensal derived metabolites have exhibited both pro- and anti-viral effects. Conversely, any impact of viral infection on microbially derived metabolites remained unknown. LC-MS/MS metabolomics has been successfully utilized to identify metabolites (32-34). Therefore, we took a similar approach to discover bacterially-derived or bacterially-dependent host-derived metabolites that are influenced by viral infection.

Abundances of 271 metabolites were found to be affected by both bacterial colonization and viral infection in the spleen (Figure 3C). The vast majority of these metabolites (250x) are unknown and consequently, their relation to known metabolites remains to be determined. Known metabolites within the sera and spleen, influenced by the combination of bacterial colonization and viral infection, were grouped by chemical type in an attempt to identify trends in the abundances of similar metabolites (Figures 3B and 3D). The only discernable trend determined by both the virus and bacterium was the increased abundance of amino acids (both essential and non-essential) in the spleen (Figure 3D).

Mechanistically how MuLV infection in *L. murinus* colonized mice alters the metabolomic landscape in the spleen remains to be determined. MuLV infection results in increased extramedullary hematopoiesis within the spleen (22), expanding the population of target cells the virus can infect. MuLV readily replicates within the proliferating erythroid progenitor cells, augmenting the chance of proviral integration near a cellular proto-oncogene and leading to the generation of pre-cancerous cells. The presence of the microbiota during the generation of these cells may alter the transcriptional landscape leading to the production of metabolites within the spleen.

Permeability of the intestinal barrier did not increase in SPF mice upon MuLV infection (Figure 4), indicating the observed increase of metabolites within the spleens of monocolonized infected mice may not be due to significant influx of metabolites originating from the gut. Thus, the origin of certain microbially dependent virally induced metabolites may be within the spleen. However, due to the diffusible nature of small gut derived metabolites we cannot rule out the gut as a source of these metabolites.

In summary, our study has demonstrated for the first time that the gut commensal bacterium and the virus together alter the metabolomic landscape of a sterile organ not directly connected with the gut.

## Acknowledgement and authors’ contribution

We thank members of the Golovkina and Chervonsky labs for helpful discussions. We also thank GRAF at University of Chicago, which enable gnotobiotic studies. This work was supported by NIH grant R01CA232882 to T.G. and M.F. J.S. was supported by T32 GM007183. This work was also supported by NIH/NIDDK Digestive Disease Research Core Center grant DK42086 and NIH grant P30CA014599.

J.S. generated mice and collected sera and spleens for metabolomic profiling. M. F. B.D. and J.M.S. performed metabolomic profiling. V. B. carried out computational analysis. M.F. and A.C. contributed to experimental design and the edit of the manuscript. J.S. and T.G. wrote the manuscript. T.G. conceived the project.

## References

1. Lee YK, Mazmanian SK. 2010. Has the microbiota played a critical role in the evolution of the adaptive immune system? Science 330:1768–73.

2. Gasaly N, de Vos P, Hermoso MA. 2021. Impact of Bacterial Metabolites on Gut Barrier Function and Host Immunity: A Focus on Bacterial Metabolism and Its Relevance for Intestinal Inflammation. Frontiers in Immunology 12.

3. Donia MS, Fischbach MA. 2015. HUMAN MICROBIOTA. Small molecules from the human microbiota. Science 349:1254766.

4. Buffie CG, Pamer EG. 2013. Microbiota-mediated colonization resistance against intestinal pathogens. Nature Reviews Immunology 13:790–801.

5. Lee W-J, Hase K. 2014. Gut microbiota–generated metabolites in animal health and disease. Nature Chemical Biology 10:416–424.

6. Sittipo P, Shim J-w, Lee YK. 2019. Microbial Metabolites Determine Host Health and the Status of Some Diseases. International Journal of Molecular Sciences. 20(21):doi:10.3390/ijms20215296.

7. Averina OV, Zorkina YA, Yunes RA, Kovtun AS, Ushakova VM, Morozova AY, Kostyuk GP, Danilenko VN, Chekhonin VP. 2020. Bacterial Metabolites of Human Gut Microbiota Correlating with Depression. International Journal of Molecular Sciences. 23(23):doi:10.3390/ijms21239234.

8. Li Z, Quan G, Jiang X, Yang Y, Ding X, Zhang D, Wang X, Hardwidge PR, Ren W, Zhu G. 2018. Effects of Metabolites Derived From Gut Microbiota and Hosts on Pathogens. Frontiers in Cellular and Infection Microbiology 8.

9. Kirby DT, Savage JM, Plotkin BJ. 2014. Menaquinone (Vitamin K2) enhancement of Staphylococcus aureus biofilm formation. Journal of Biosciences and Medicines 2014.

10. Yang Y, Yao F, Zhou M, Zhu J, Zhang X, Bao W, Wu S, Hardwidge PR, Zhu G. 2013. F18ab Escherichia coli flagella expression is regulated by acyl-homoserine lactone and contributes to bacterial virulence. Veterinary Microbiology 165:378–383.

11. Hochbaum Allon I, Kolodkin-Gal I, Foulston L, Kolter R, Aizenberg J, Losick R. 2011. Inhibitory Effects of d-Amino Acids on Staphylococcus aureus Biofilm Development. Journal of Bacteriology 193:5616–5622.

12. Araki S, Suzuki M, Fujimoto M, Kimura M. 1995. Enhancement of Resistance to Bacterial Infection in Mice by Vitamin B2. Journal of Veterinary Medical Science 57:599–602.

13. Jacobson A, Lam L, Rajendram M, Tamburini F, Honeycutt J, Pham T, Van Treuren W, Pruss K, Stabler SR, Lugo K, Bouley DM, Vilches-Moure JG, Smith M, Sonnenburg JL, Bhatt AS, Huang KC, Monack D. 2018. A Gut Commensal-Produced Metabolite Mediates Colonization Resistance to Salmonella Infection. Cell Host & Microbe 24:296-307.e7.

14. Rowland I, Gibson G, Heinken A, Scott K, Swann J, Thiele I, Tuohy K. 2018. Gut microbiota functions: metabolism of nutrients and other food components. European Journal of Nutrition 57:1–24.

15. Sánchez-García FJ, Pérez-Hernández CA, Rodríguez-Murillo M, Moreno-Altamirano MMB. 2021. The Role of Tricarboxylic Acid Cycle Metabolites in Viral Infections. Frontiers in Cellular and Infection Microbiology 11.

16. Fontaine Krystal A, Sanchez Erica L, Camarda R, Lagunoff M. 2015. Dengue Virus Induces and Requires Glycolysis for Optimal Replication. Journal of Virology 89:2358–2366.

17. Alwin A, Karst SM. 2021. The influence of microbiota-derived metabolites on viral infections. Curr Opin Virol 49:151–156.

18. Trompette A, Gollwitzer ES, Pattaroni C, Lopez-Mejia IC, Riva E, Pernot J, Ubags N, Fajas L, Nicod LP, Marsland BJ. 2018. Dietary Fiber Confers Protection against Flu by Shaping Ly6c− Patrolling Monocyte Hematopoiesis and CD8+ T Cell Metabolism. Immunity 48:992-1005.e8.

19. Spring J, Khan AA, Lara S, O’Grady K, Wilks J, Gurbuxani S, Erickson S, Fischbach M, Jacobson A, Chervonsky A, Golovkina T. 2022. Gut commensal bacteria enhance pathogenesis of a tumorigenic murine retrovirus. Cell Reports 40.

20. Wilks J, Beilinson H, Theriault B, Chervonsky A, Golovkina T. 2014. Antibody-mediated immune control of a retrovirus does not require the microbiota. J Virol 88:6524–7.

21. Sarma-Rupavtarm RB, Ge Z, Schauer DB, Fox JG, Polz MF. 2004. Spatial distribution and stability of the eight microbial species of the altered schaedler flora in the mouse gastrointestinal tract. Appl Environ Microbiol 70:2791–800.

22. Hook LM, Jude BA, Ter-Grigorov VS, Hartley JW, Morse HC, 3rd, Trainin Z, Toder V, Chervonsky AV, Golovkina TV. 2002. Characterization of a novel murine retrovirus mixture that facilitates hematopoiesis. J Virol 76:12112–22.

23. Rowe WP, Pugh WE, Hartley JW. 1970. Plaque assay techniques for murine leukemia viruses. Virology 42:1136–1139.

24. Volynets V, Reichold A, Bárdos G, Rings A, Bleich A, Bischoff SC. 2016. Assessment of the Intestinal Barrier with Five Different Permeability Tests in Healthy C57BL/6J and BALB/cJ Mice. Digestive Diseases and Sciences 61:737–746.

25. Noecker C, Sanchez J, Bisanz JE, Escalante V, Alexander M, Trepka K, Heinken A, Liu Y, Dodd D, Thiele I, DeFelice B, Turnbaugh PJ. 2022. Systems biology illuminates alternative metabolic niches in the human gut microbiome. bioRxiv doi:10.1101/2022.09.19.508335:2022.09.19.508335.

26. Brenchley JM, Price DA, Schacker TW, Asher TE, Silvestri G, Rao S, Kazzaz Z, Bornstein E, Lambotte O, Altmann D, Blazar BR, Rodriguez B, Teixeira-Johnson L, Landay A, Martin JN, Hecht FM, Picker LJ, Lederman MM, Deeks SG, Douek DC. 2006. Microbial translocation is a cause of systemic immune activation in chronic HIV infection. Nat Med 12:1365–71.

27. Labarta-Bajo L, Nilsen SP, Humphrey G, Schwartz T, Sanders K, Swafford A, Knight R, Turner JR, Zúñiga EI. 2020. Type I IFNs and CD8 T cells increase intestinal barrier permeability after chronic viral infection. Journal of Experimental Medicine 217.

28. Baldridge MT, Nice TJ, McCune BT, Yokoyama CC, Kambal A, Wheadon M, Diamond MS, Ivanova Y, Artyomov M, Virgin HW. 2015. Commensal microbes and interferon-lambda determine persistence of enteric murine norovirus infection. Science 347:266–9.

29. Kane M, Case LK, Kopaskie K, Kozlova A, MacDearmid C, Chervonsky AV, Golovkina TV. 2011. Successful transmission of a retrovirus depends on the commensal microbiota. Science 334:245–9.

30. Kuss SK, Best GT, Etheredge CA, Pruijssers AJ, Frierson JM, Hooper LV, Dermody TS, Pfeiffer JK. 2011. Intestinal microbiota promote enteric virus replication and systemic pathogenesis. Science 334:249–252.

31. Uchiyama R, Chassaing B, Zhang B, Gewirtz AT. 2014. Antibiotic treatment suppresses rotavirus infection and enhances specific humoral immunity. J Infect Dis 210:171–82.

32. Yoshimoto S, Loo TM, Atarashi K, Kanda H, Sato S, Oyadomari S, Iwakura Y, Oshima K, Morita H, Hattori M, Honda K, Ishikawa Y, Hara E, Ohtani N. 2013. Obesity-induced gut microbial metabolite promotes liver cancer through senescence secretome. Nature 499:97–101.

33. Sharon G, Cruz NJ, Kang D-W, Gandal MJ, Wang B, Kim Y-M, Zink EM, Casey CP, Taylor BC, Lane CJ, Bramer LM, Isern NG, Hoyt DW, Noecker C, Sweredoski MJ, Moradian A, Borenstein E, Jansson JK, Knight R, Metz TO, Lois C, Geschwind DH, Krajmalnik-Brown R, Mazmanian SK. 2019. Human Gut Microbiota from Autism Spectrum Disorder Promote Behavioral Symptoms in Mice. Cell 177:1600-1618.e17.

34. Hsiao EY, McBride SW, Hsien S, Sharon G, Hyde ER, McCue T, Codelli JA, Chow J, Reisman SE, Petrosino JF, Patterson PH, Mazmanian SK. 2013. Microbiota modulate behavioral and physiological abnormalities associated with neurodevelopmental disorders. Cell 155:1451–1463.

35. Chen SY, Feng Z, Yi X. 2017. A general introduction to adjustment for multiple comparisons. J Thorac Dis 9:1725–1729.

