## Supplemental Table 1 for "Retroviral infection and commensal bacteria dependently alter the metabolomic profile in a sterile organ"

| 5-Methylcytidine | Creatine | Isoleucine | Creatinine | Leucine | Acetylcarnitine | Cystathionine | Linoleoylcarnitine | Alanine |
| --- | --- | --- | --- | --- | --- | --- | --- | --- |
| Cytidine | Lysine | Methionine | Nicotinamide | Riboflavin | Betaine | Glu-Thr | 2'-Deoxycytidine | Carnitine |
| 3-Methylcytidine | Choline [M]+ | homocitrulline | Taurine | Hypotaurine | Pipecolic acid | Phenylalanine | Urea | Arginine |
| Valine | Ornithine | Proline | Citrulline | Oleoyl-L-carnitine | Serine | N,N-Dimethylarginine | Guanidinoacetic acid | Palmitoylcarnitine |
| 7,8-Dihydrobiopterin | gamma-Glutamylleucine | N6,N6-dimethyllysine | 1-Methyladenosine | N-epsilon-Acetyllysine | N-.alpha.-Acetyl-L-arginine | Palmitoyl sphingomyelin | Myristoyl-L-carnitine | Methioninesulfoxide [M+H]+ |
| 2-Arachidonoylglycerol | Nalpha-acetyl-L-Lysine | SN-Glycero-3-Phosphocoline | 1-Palmitoyl-sn-glycero-3-phosphocholine late | 5-Methyl-2'-deoxycytidine | N-Oleoyl-D-erythro-sphingosylphosphorylcholine | Palmitoyleicosapentaenoyl phosphatidylcholine | 1-Stearoyl-2-hydroxy-sn-glycero-3-phosphocholine | 1-Heptadecanoyl-sn-glycero-3-phosphocholine late |
| 1,2-Dilinoleoyl-sn-glycero-3-phosphocholine | 1-(1Z-Octadecenyl)-2-(5Z,8Z,11Z,14Z-eicosatetraenoyl)-sn-glycero-3-phosphocholine | 1-Oleoyl-sn-glycero-3-phosphoethanolamine | 1-(1Z-Octadecenyl)-2-(4Z,7Z,10Z,13Z,16Z,19Z-docosahexaenoyl)-sn-glycero-3-phosphocholine | 1-O-Hexadecyl-2-O-(5Z,8Z,11Z,14Z,17Z-eicosapentaenoyl)-sn-glyceryl-3-phosphorylcholine | 1-(1Z-Hexadecenyl)-sn-glycero-3-phosphocholine late | 1-O-Hexadecyl-2-O-(4Z,7Z,10Z,13Z,16Z,19Z-docosahexaenoyl)-sn-glyceryl-3-phosphorylcholine | 1-Stearoyl-2-hydroxy-sn-glycero-3-phosphoethanolamine | 1-Hexadecyl-2-(5Z,8Z,11Z,14Z-eicosatetraenoyl)-sn-glycero-3-phosphocholine |
| 1,2-dioleoyl-sn-glycero-3-phosphatidylcholine | 1-(1Z-Octadecenyl)-sn-glycero-3-phosphocholine early | 1-Palmitoyl-2-docosahexaenoyl-sn-glycero-3-phosphocholine | 1-(1Z-Hexadecenyl)-sn-glycero-3-phosphocholine early | 1-Myristoyl-sn-glycero-3-phosphocholine | 1-O-Octadecyl-sn-glyceryl-3-phosphorylcholine | 1-O-Hexadecyl-2-O-acetyl-sn-glyceryl-3-phosphorylcholine | 1-(1Z-Octadecenyl)-sn-glycero-3-phosphocholine late | 1-Palmitoyl-sn-glycero-3-phosphocholine early |
| 1,2-Dipalmitoleoyl-sn-glycero-3-phosphocholine |  |  |  |  |  |  |  |  |

Table S1: Known metabolites in the sera under positive ionization. Metabolites identified in the sera of GF, GF infected, *L. murinus* colonized, and *L. murinus* colonized and infected mice under positive ionization.
