## Supplemental Table 2 for "Retroviral infection and commensal bacteria dependently alter the metabolomic profile in a sterile organ"

| Citrulline | 4-Nitrophenol | Cytidine | Allantoin | Hypotaurine | Arabitol | Isoleucine | Adipic acid |
| --- | --- | --- | --- | --- | --- | --- | --- |
| Arginine | Leucine | Asparagine | lysoPC 14:0 | Citric acid | Vanillin | Thymidine | Taurine |
| Serine | Pseudouridine | myo-Inositol | lysoPC 16:0 early | lysoPC 18:2 late | lysoPC 18:0 late | lysoPE 16:0 late | lysoPC 18:2 early |
| Delta-Hydroxylysine | lysoPC 18:0 early | lysoPE 16:0 early | lysoPC 17:0 late | lysoPC 20:4 late | lysoPC 16:0 late | lysoPC 20:4 early | N-Acetylalanine |
| lysoPC 18:1 late | lysoPE 18:0 late | lysoPC 18:1 early | lysoPE 18:0 early | Pantothenic acid | Methioninesulfoxide | N-epsilon-Acetyllysine | N-Acetylleucine |
| Palmitoyl sphingomyelin | N-Acetylhistidine | N,N-Dimethylarginine | N-Methylglutamic acid | N-.alpha.-Acetyl-L-ornithine | Methyl-beta-galactopyranoside | 2-Deoxyuridine | 2-Mercaptobenzothiazole |
| 2-Hydroxypalmitic acid | Beta.-Glycerophosphate | 3-Hydroxyoctanoic acid | 1-Palmitoyl-2-linoleoyl-sn-glycero-3-phosphocholine | 1-Palmitoyl-2-docosahexaenoyl-sn-glycero-3-phosphocholine | 1-Palmitoyl-2-arachidonoyl-sn-glycero-3-phosphocholine | 1-(1Z-Hexadecenyl)-sn-glycero-3-phosphocholine |  |

Table 2: Metabolites identified in the sera under negative ionization. Known metabolites in the sera of GF, GF infected, *L. murinus* colonized, and *L. murinus* colonized plus infected mice under negative ionization.
