## Supplemental Table 3 for "Retroviral infection and commensal bacteria dependently alter the metabolomic profile in a sterile organ"

| Nicotinamide | Ethanolamine | Anserine | Tyrosine | Pyridoxal | Threonine | Valine | Carnitine |
| --- | --- | --- | --- | --- | --- | --- | --- |
| Hypoxanthine | Pyrazinamide | Taurine | Urea | Tryptophan | Cytidine | Arginine | Phenylalanine |
| Aspartic acid | Methionine | Uracil | Xanthine | Choline [M]+ | Serine | Proline | Alanine |
| Glutamic acid | Asparagine | Histidine | Thiamine Cation | Argininosuccinic acid | Sphinganine | Glucose-6-phosphate | N-epsilon-Acetyllysine |
| 2'-Deoxycytidine | N,N-Dimethylarginine | Ergothioneine | 3'-O-Methylinosine | Epsilon.-Caprolactam | N-Acetyl-D-glucosamine | Palmitoyl sphingomyelin | Nicotinamide riboside cation |
| N.epsilon.-Methyl-L-lysine | Gamma.-Aminobutyric acid | Methioninesulfoxide [M+H]+ | 1,2-Cyclohexanedione | 1,2-Dilinoleoyl-sn-glycero-3-phosphocholine | 1-Myristoyl-2-stearoyl-sn-glycero-3-phosphocholine | (2R)-3-Hydroxyisovaleroylcarnitine | 1,2-Diarachidonoyl-sn-glycero-3-phosphocholine |
| 1-O-Hexadecyl-2-O-(4Z,7Z,10Z,13Z,16Z,19Z-docosahexaenoyl)-sn-glyceryl-3-phosphorylcholine | 1-(1Z-Octadecenyl)-2-(5Z,8Z,11Z,14Z-eicosatetraenoyl)-sn-glycero-3-phosphoethanolamine | 1-(1Z-Octadecenyl)-2-(4Z,7Z,10Z,13Z,16Z,19Z-docosahexaenoyl)-sn-glycero-3-phosphoethanolamine | 2-Docosahexaenoyl-1-stearoyl-sn-glycero-3-phosphoethanolamine | 1-Stearoyl-2-linoleoyl-sn-glycero-3-phospho-L-serine | 1,2-Dipentadecanoyl-sn-glycero-3-phosphocholine | 2-Arachidonoyl-1-palmitoyl-sn-glycero-3-phosphoethanolamine | 1,2-dioleoyl-sn-glycero-3-phosphatidylcholine |
| 1-Palmitoyl-2-docosahexaenoyl-sn-glycero-3-phosphocholine | 1-Oleoyl-2-myristoyl-sn-glycero-3-phosphocholine | 2-Oleoyl-1-palmitoyl-sn-glycero-3-phosphocholine | Nepsilon, Nepsilon-Trimethyllysine |  |  |  |  |

Table 3: Known metabolites found in spleens under positive ionization. Metabolites identified in the spleens of GF, GF infected, *L. murinus* colonized, and *L. murinus* colonized and infected mice under positive ionization.
