## Supplemental Table 4 for "Retroviral infection and commensal bacteria dependently alter the metabolomic profile in a sterile organ"

| PE 38:4 | Galactitol | Uracil | Xanthine | Arginine | Gly-Asp | Taurine | myo-Inositol |
| --- | --- | --- | --- | --- | --- | --- | --- |
| Threonine | Hypoxanthine | N-Acetylserine | 2'-Deoxycytidine | Xylitol | N-Acetylalanine | 2-Deoxyuridine | Allantoin |
| Aspartic acid | Glucose | PC 32:0 | Serine | Threonic acid | PE 38:6 | PC 32:1 | Glutamine |
| D-Asparagine | Histidine | Gly-Gly | PS 36:2 | PE 36:4 | Valine | Orotic acid | Glutamic acid |
| Ribose-5-phosphate | Tyrosine | Cytidine | Pseudouridine | N-epsilon-Acetyllysine | Glucose-1-phosphate | Dihydroxyacetone phosphate | Glucose-6-phosphate |
| N,N-Dimethylarginine | Pyroglutamic acid | Palmitoyl sphingomyelin | Ribose-1-phosphate | Plasmenyl-PE 38:6 | L-.gamma.-Glutamyl-L-glutamic acid | Plasmenyl-PE 36:4 | Plasmenyl-PE 38:4 |
| O-Phosphocolamine | 2-Docosahexaenoyl-1-stearoyl-sn-glycero-3-phosphoserine | 1-Stearoyl-2-arachidonoyl-sn-glycero-3-phosphoserine | 1-Palmitoyl-2-docosahexaenoyl-sn-glycero-3-phosphocholine |  |  |  |  |

Table 4: Identified metabolites found in spleens under negative ionization. Known metabolites identified in the spleens of GF, GF infected, *L. murinus* colonized, and *L. murinus* colonized and infected mice under negative ionization.
